# The D614G mutation in the SARS-CoV-2 spike protein reduces S1 shedding and increases infectivity

**DOI:** 10.1101/2020.06.12.148726

**Authors:** Lizhou Zhang, Cody B Jackson, Huihui Mou, Amrita Ojha, Erumbi S Rangarajan, Tina Izard, Michael Farzan, Hyeryun Choe

## Abstract

SARS coronavirus 2 (SARS-CoV-2) isolates encoding a D614G mutation in the viral spike (S) protein predominate over time in locales where it is found, implying that this change enhances viral transmission. We therefore compared the functional properties of the S proteins with aspartic acid (S^D614^) and glycine (S^G614^) at residue 614. We observed that retroviruses pseudotyped with S^G614^ infected ACE2-expressing cells markedly more efficiently than those with S^D614^. This greater infectivity was correlated with less S1 shedding and greater incorporation of the S protein into the pseudovirion. Similar results were obtained using the virus-like particles produced with SARS-CoV-2 M, N, E, and S proteins. However, S^G614^ did not bind ACE2 more efficiently than S^D614^, and the pseudoviruses containing these S proteins were neutralized with comparable efficiencies by convalescent plasma. These results show S^G614^ is more stable than S^D614^, consistent with epidemiological data suggesting that viruses with S^G614^ transmit more efficiently.

---

Until late 2019, only six coronaviruses were known to infect humans: HCoV-229E, HCoV-OC43, SARS-CoV (SARS-CoV-1), HCoV-NL63, CoV-HKU1, and MERS-CoV. A seventh, SARS-CoV-2, emerged in the winter of 2019 from Wuhan, China. SARS-CoV-2 is closely related to SARS-CoV-1, a virus that appeared from Guangdong province, China in late 2002.

The coronavirus spike (S) protein mediates receptor binding and fusion of the viral and cellular membrane. The S protein extends from the viral membrane and is uniformly arranged as trimers on the virion surface to give the appearance of a crown (*corona* in Latin). The coronavirus S protein is divided into two domains: S1 and S2. The S1 domain mediates receptor binding, and the S2 mediates downstream membrane fusion^1,2^. The receptor for SARS-CoV-2 is angiotensin-converting enzyme 2 (ACE2)^3–7^, a metalloprotease that also serves as the receptor for SARS-CoV-1^8^. A small, independently folded subdomain of S1, described as the receptor-binding domain (RBD), directly binds ACE2 when the virus engages a target cell^9–12^. The S1/S2 junction of SARS-CoV-2 is processed by a furin-like proprotein convertase in the virus producer cell. In contrast, the S1/S2 junction of SARS-CoV-1 is processed by TMPRSS2 at the cell surface or by lysosomal cathepsins in the target cells^13–18^. Both S proteins are further processed in the target cell within the S2 domain at the S2’ site, an event that is also required for productive infection^19,20^.

Recent analyses of the fine-scale sequence variation of SARS-CoV-2 isolates identified several genomic regions of increased genetic variation^21–30^. One of these variations encodes a S-protein mutation, D614G, in the carboxy(C)-terminal region of the S1 domain^21–23,26,30^. This region of the S1 domain directly associates with S2 (Fig. 1a). This mutation with glycine at the residue 614 (G614) was previously detected to increase with an alarming speed^21,22^. Our own analysis of the S-protein sequences available from the GenBank showed a similar result: The G614 genotype was not detected in February (among 33 sequences) and observed at low frequency in March (26%), but increased rapidly by April (65%) and May (70%) (Fig. 1b), indicating a transmission advantage over viruses with D614. Korber et al. noted that this change also correlated with increased viral loads in COVID-19 patients^22^, but because this change is also associated with the mutations in viral nsp3 and RdRp proteins, the role of the S-protein in these observations remained undefined.

**Figure 1.**
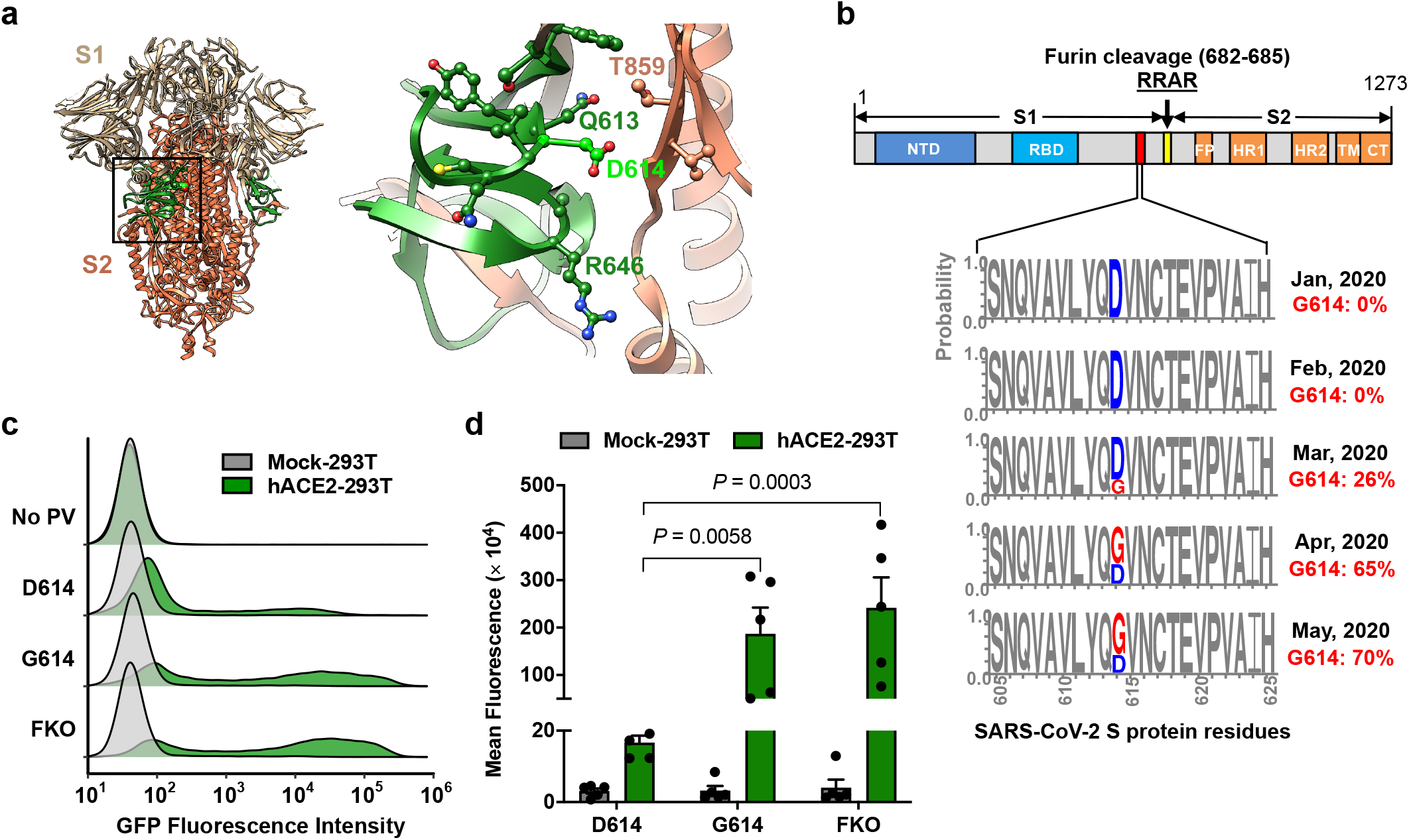
The D614G mutation is associated with enhanced infectivity. Cryo-EM structure of S1 (grey) and S2 (orange) heterodimer (PBD 6VXX). The residues 581-676, a C-terminal region of the S1 domain involved in S2 interaction, is shown in green. Aspartic acid 614 is shown in light green. The area indicated with a black square is presented magnified at the right. Residues within 5.5 Å of D614 are shown in a ball-and-stick representation. **b,** A representation of the SARS-CoV-2 Sprotein (upper panel) and D/G variation at the residue 614 presented in logo plots at different time points between January 1^st^ and May 30^th^, 2020 (lower panel). Total number of sequences analyzed: 17 in January, 33 in February, 293 in March, 1511 in April, and 2544 in May. NTD: N-terminal domain, RBD: Receptor-binding domain, FP: Fusion peptide, HR1 and HR2: Heptad-repeat region 1 and 2, respectively, TM: Transmembrane region, CT: Cytoplasmic tail. **c,d,** Mock-and hACE2-293T cells on 96-well plates were infected with MLV PV (5 x 10^8^ vector genome per well) expressing GFP and pseudotyped with the indicated viral glycoprotein and analyzed 24 h later. Representative histograms (c) or mean ± SEM (d) of five experiments conducted using two independent PV preparations are shown. Each dot in (d) indicates an average value of a duplicated experiment. Significant differences were analyzed by two-way ANOVA with Sidak multiple comparisons test. PV titers are presented in Extended Data Fig. 1. FKO: Furincleavage knockout mutant.

To determine if the D614G mutation alters the properties of the S-protein in a way that could impact transmission or replication, we assessed its role in viral entry. Maloney murine leukemia virus (MLV)-based pseudoviruses (PVs), expressing green fluorescent protein (GFP) and pseudotyped with the S protein of SARS-CoV-2 (SARS2) carrying the D614 or G614 genotype (S^D614^ and S^G614^, respectively) were produced from transfected HEK293T cells as previously described^31^. An S^D614^ variant, in which the furin-cleavage motif between the S1 and S2 domains is ablated (S^FKO^), was also included for comparison. HEK293T cells transduced to express human ACE2 (hACE2-293T) or those transduced with vector alone (Mock-293T) were infected with the same particle numbers of the PVs pseudotyped with the S^D614^, S^G614^, or S^FKO^ (PV^D614^, PV^G614^, or PV^FKO^, respectively), and infection level was assessed one day later. We observed PV^G614^ infected hACE2-293T cells with approximately 9-fold higher efficiency than did PV^D614^ (Fig. 1c,d). This enhanced infectivity of PV^G614^ is not an artifact of PV titer normalization, as their titers are very similar (Extended Data Fig. 1).

We next investigated the mechanism with which S^G614^ increased virus infectivity. Because S1 residue 614 is proximal to the S2 domain, we first compared the ratio between the S1 and S2 domains in the virion that might indicate altered release or shedding of the S1 domain after cleavage at the S1/S2 junction. To do so, we used S-protein constructs bearing Flag tags at both their amino (N)- and C-termini. PVs pseudotyped with these double-Flag tagged forms of S^D614^, S^G614^, and S^FKO^ were partially purified and concentrated by pelleting through a 20% sucrose layer^32^ and evaluated for their infectivity. The titers of PVs were similar among PV^G614^, PV^D614^, and PV^FKO^ before and after purification (Fig. 2a). In addition, modification by Flag-tags or pelleting of PVs through a sucrose layer did not alter the relative infectivity between PV^G614^ and PV^D614^ (Fig. 2b). We then determined their S1:S2 ratio by western blotting using the anti-Flag M2 antibody. As shown in Fig. 2c, the S1:S2 ratio is markedly greater in PV^G614^ compared to PV^D614^, indicating that glycine at residue 614 of S^G614^ stabilizes the interaction between the S1 and S2 domains, limiting S1 shedding. In addition, the total amount of the S protein in PV^G614^ is also much higher than that in PV^D614^, as indicated by a denser S2 band, even though the same number of pseudovirions was analyzed, as determined by quantitative PCR. To independently confirm that similar number of virions was analyzed, the lower part of the same membrane was blotted with an anti-p30 MLV gag antibody (Fig. 2c). Similar densities of p30 bands were observed from all PVs, indicating that differences in S-protein incorporation observed with PV^G614^ and PV^D614^ were due to the mutation of residue 614, not by different amount of PVs analyzed. A similar experiment performed with independently produced PVs yielded a nearly identical result (Extended Data Fig. 2a). Densitometric analysis shows there is 4.7 times more S1+S2 band in PV^G614^ compared to PV^D614^ (Fig. 2d). To more accurately estimate the S1:S2 ratio, we next compared different amount of the same samples so that S2-band intensity in PV^G614^ and PV^D614^ was comparable (Fig. 2e). Averages of several quantification show that the S1:S2 ratio of PV^G614^ is 3.5 times higher than that of PV^D614^ (Fig. 2f). The M2 antibody used in this experiment binds the Flag tag located at both the N- and C-termini of a protein, but it binds N-terminal Flag tag more efficiently^33^. Therefore, we directly visualized virion S-protein bands by silver staining (Extended Data Fig. 2b). Although the S2 bands are masked by a same-sized MLV protein, S1 bands are well separated. Again, the intensity of the S1 band of PV^G614^ is much stronger than that of PV^D614^, while p30 bands are comparable, a result consistent with those observed using the anti-Flag M2 antibody.

**Figure 2.**
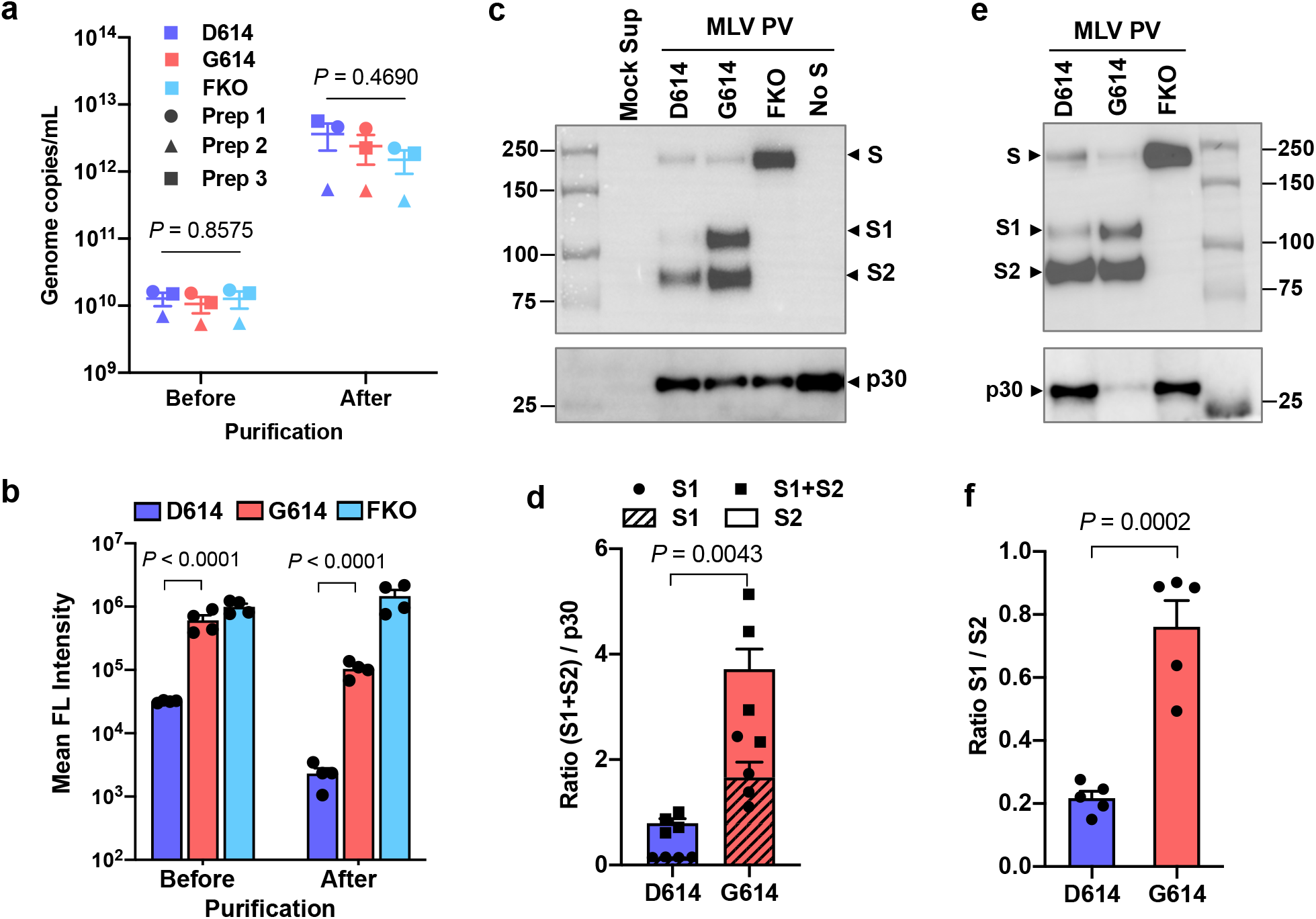
Superior infectivity of G614 results from decreased S1 shedding and higher level of S protein in the virion. **a-f,** Indicated MLV PVs produced with the S protein containing the Flag tag at both the N-and C-termini were partially purified and concentrated by pelleting through a 20% sucrose layer. PV titers were assessed by RT-qPCR (a). The same symbols in different PV groups indicate they are from the same batch. The same PVs were assessed for their infectivity in Mock- and hACE2-293T cells (b). Each symbol in (a,b) indicates an average value of a duplicated experiment. Mean ± SEM of three independently prepared PVs (a) and four experiments using those three PV batches (b) are shown. The same amount (1 x 10^10^ vg per lane) (c,d) or to more accurately compare the S1 and S2 ratio, different amount (e) of the purified PVs were analyzed by WB using the anti-Flag M2 antibody or anti-p30 MLV gag antibody. A similar experiment performed with an independently prepared batch of PVs is shown in Extended Data Fig 2a. The same PVs visualized by silver stain is shown in Extended Data Fig. 2b. Total virion S protein (d) and the S1/S2 ratio (f) of PVD^614^ and PV^G614^ were calculated from four (d) or five (f) WBs performed with three independently prepared PV batches and presented as mean ± SEM. Significant differences were analyzed by one-way ANOVA (a), two-way ANOVA with Sidak multiple comparison tests of log-transformed data (b), or unpaired Student’s t-test (d,f).

We next confirmed these findings using virus-like particles (VLPs) composed only of the native SARS-CoV-2 proteins, the nucleoprotein (N), membrane protein (M), envelope protein (E), and S protein^34^. VLPs were partially purified and analyzed in the same way as MLV PVs. The S protein bands were detected with the anti-Flag M2 antibody, and the N protein with pooled convalescent plasma derived from COVID-19 patients. The S1:S2 ratio and total S protein on the virion was again much higher in the VLPs carrying S^G614^ (VLP^G614^) compared to those carrying S^D614^ (VLPD^614^) (Fig. 3a). The S1:S2 ratio is 3.4 fold higher and the total S protein is nearly five fold enriched in VLP^G614^ compared to VLPD^614^ (Fig. 3b,c). Thus, the D614G mutation enhances virus infection through two related mechanisms: It reduces S1 shedding and increases total S protein incorporated into the virion.

**Fig. 3.**
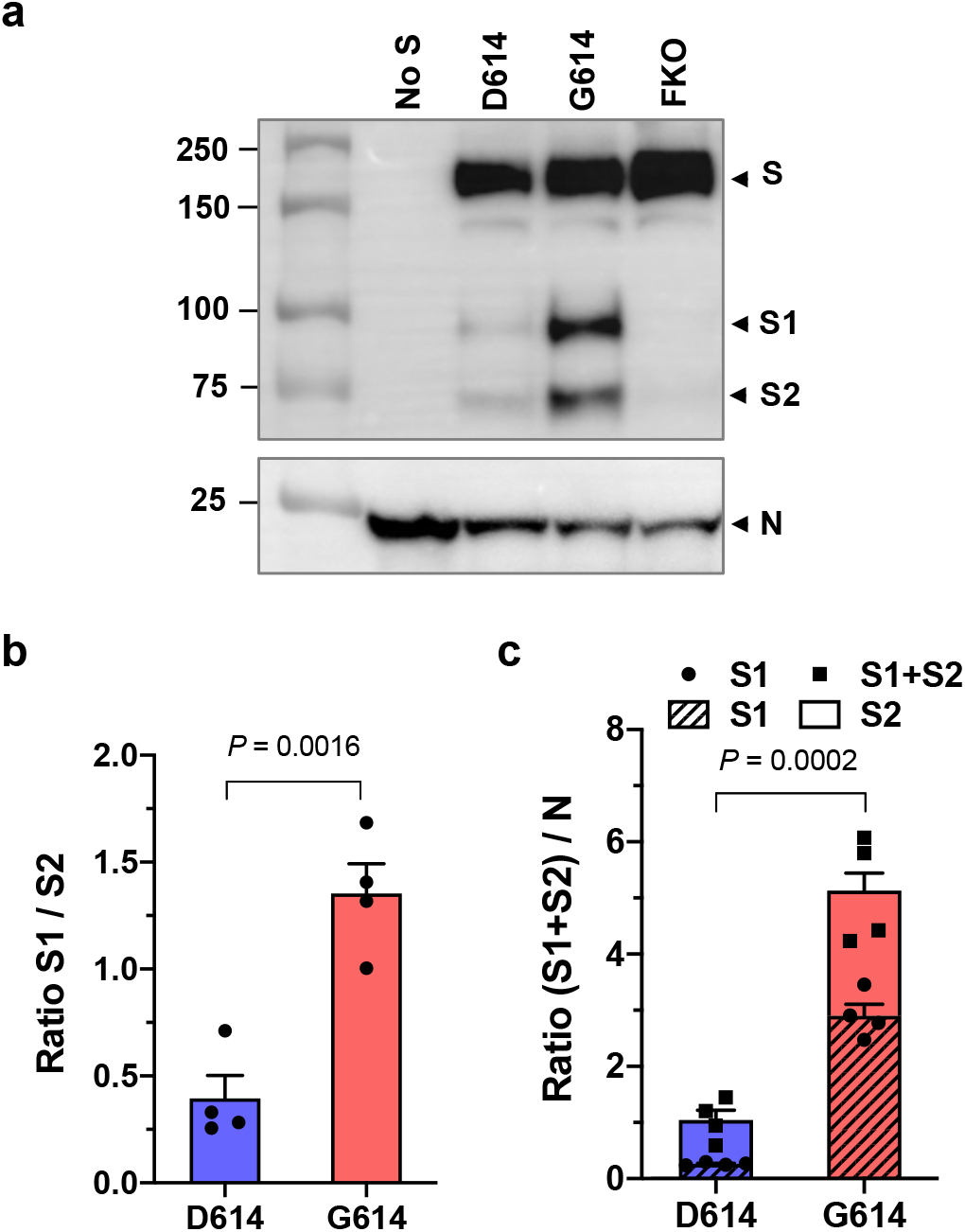
SARS-CoV-2 VLP^G614^ also exhibits decreased S1 shedding and increased total virion S protein. **a-c,** VLPs were produced from HEK293T cell transfection of the M, N, E, and S proteins of SARS-CoV-2. VLPs were harvested from the culture supernatant and partially purified as MLV PVs. The S protein bands were visualized using the anti-Flag tag M2 antibody and the N protein band using pooled convalescent plasma (a). A representative of WBs performed with three independently prepared VLPs is shown. The S1/S2 ratio (b) and the difference in total virion S protein (c) incorporated into the VLPD^614^ and VLP^G614^ were calculated from four WBs performed with three independent VLP preparations and presented as mean ± SEM. Significant differences were analyzed by unpaired Student’s t-test.

It has previously been speculated that D614G mutation promotes an open configuration of the S protein that is more favorable to ACE2 association^5,22,23,35^. To explore this possibility, we investigated whether ACE2 binding by S^G614^ was more efficient than that by S^D614^. HEK293T cells transfected to express each S protein were assessed for their binding of hACE2 immunoadhesin, using hACE2-NN-Ig, whose enzymatic activity was abolished by mutation^31^. hACE2-NN-Ig bound SARS-CoV-1 RBD with an equivalent affinity as hACE2-Ig. These S proteins are fused to a C-terminal, but not an N-terminal Flag tag, thus allowing for the measurement of total S protein expression in permeabilized cells by flow cytometry. Although total S protein expression was comparable, hACE2-NN-Ig binding to the cells expressing S^G614^ was substantially higher than its binding to cells expressing S^D614^ (Fig. 4a). This observation has several explanations. First, G614 could increase hACE2 association by promoting greater exposure of the RBD, or second, this mutation could increase the number of binding sites by limiting S1-domain shedding. To differentiate these possibilities, we appended the Myc-tag to the N-terminus of the S-protein that is Flag tagged at its C-terminus and repeated the study, this time detecting the S1 domain with an anti-Myc antibody. As shown in Fig. 4b, the ratio of Myc-tag to Flag-tag is higher in cells expressing S^G614^ than in cells expressing S^D614^. However, the Myc-tag/Flag-tag ratio is similar to the hACE2-NN-Ig/Flag-tag ratio, indicating that increased hACE2 binding to the S^G614^-expressing cells did not result from increased affinity of S^G614^ spikes to hACE2 or greater access to the RBD. Instead, these data show there is more S1 domain in the S^G614^-expressing cells, a result again consistent with the observation that the D614G mutation reduces S1 shedding. We then assessed whether differential amount of the S protein could influence neutralization sensitivity of the virus. Fig. 4c shows that PV^D614^ and PV^G614^ are similarly susceptible to neutralizing antisera, indicating that antibody-mediated control of viruses carrying S^D614^ and S^G614^ would be similar.

**Fig 4.**
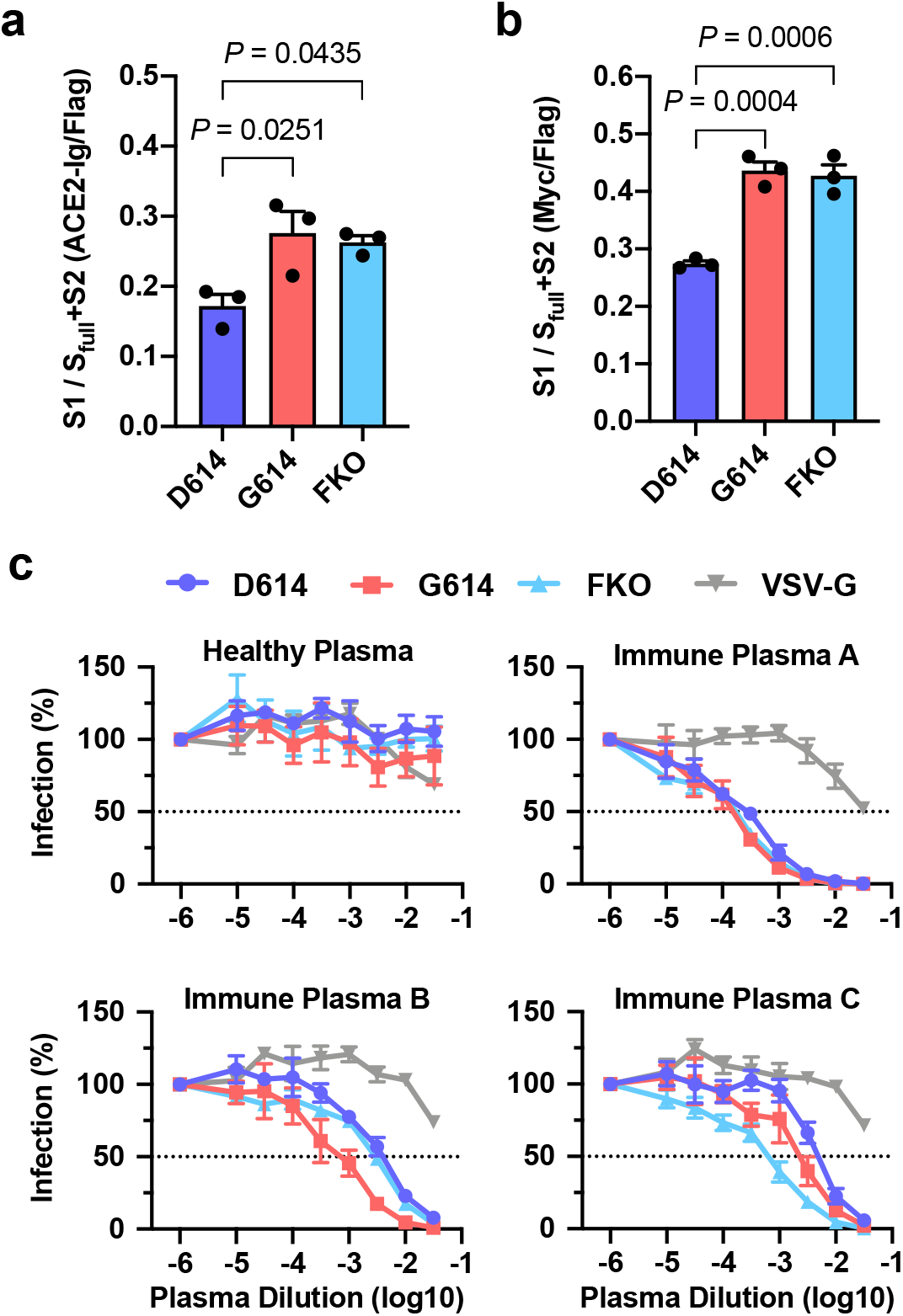
PV^G614^ is not more resistant to neutralization than does PV^D614^. **a,** The S protein containing C-terminal Flag tag is transfected into HEK293T cells and assessed for hACE2-NN-Ig binding. Total S protein was measured by detecting the Flag tag in the permeabilized cells. The ratio of hACE2-NN-Ig binding to Flag-tag staining is shown. **b,** Experiments similar to those in (a) except the S protein contains N-Myc and C-Flag tags, and S1 level was assessed using an anti-Myc antibody. Each symbol in (a,b) indicates an average value of a duplicated experiment. The data in (a,b) before normalization are presented in Extended Data Fig. 3a,b. Mean ± SEM of three independent experiments are presented. Significant differences were analyzed by one-way ANOVA and Sidak multiple comparisons test. **c,** MLV PVs expressing firefly luciferase and pseudotyped with the indicated S protein or VSV G protein were preincubated without (presented at x = −6) or with serially diluted plasmas derived from convalescent COVID-19 patients or a SARS-CoV naïve individual. hACE2-293T cells were infected with these preincubated mixes and infection was assessed 24 h later by measuring luciferase activity. Mean ± SEM of three-five independent experiments are presented.

It has also been speculated that the D614G mutation would promote, not limit, shedding of the S1 domain, based on the hypothetical loss of a hydrogen bond between D614 in S1 and T859 in S2^22^. An alternative explanation, more consistent with the data presented here, is that Q613 forms a hydrogen bond with T859, and the greater backbone flexibility provided by introduction of glycine at an adjacent position 614 enables a more favorable orientation of Q613. It is also possible that D614 can form an intra-domain salt bridge with R646, promoting a local S1 conformation unfavorable to its association with S2. In this model, replacing aspartic acid with glycine at the position 614 would prevent sampling of this unfavorable configuration. The instability of S^D614^ may also account for the observed lower level of incorporation of the functional S protein into PVs and VLPs. Specifically, the S-protein trimers with the exposed S2 domains, as a result of S1 shedding, could destabilize the trans-Golgi network membrane, the site of processing of the S1/S2 boundary, and such disruption may impede S-protein incorporation into the virion. In case of VLPs, this disruption would presumably further interfere with appropriate M- and N-protein associations by altering the conformation and orientation of the S-protein membrane-proximal regions. Alternatively, these S-protein trimers with the exposed S2 domains may serve as poor substrates for downstream post-translational modifications including palmitoylation, and those lacking proper modifications might be unsuitable for virion incorporation.

An interesting question is why viruses carrying the more stable S^G614^ appear to be more transmissible without resulting in a major observable difference in disease severity^22,27^. It is possible that higher levels of functional S protein observed with S^G614^ increase the chance of host-to-host transmission, but that other factors limit the rate and efficiency of intra-host replication. Alternatively, the loss of virion-associated S proteins observed with S^D614^ may be compensated by greater fusion efficiency with the destabilized S protein when the next target cell is adjacent in a host tissue. It is also possible that our ability to detect sequence changes at this early stage of the pandemic is simply greater than our ability to detect modest differences in pathogenesis. The strong phenotypic difference we observe here between D614 and G614 suggests that more study on the impact of the D614G mutation on the course of disease is warranted.

Finally, our data raise interesting questions about the natural history of SARS-CoV-2 as it moved presumably from horseshoe bats to humans. At some point in this process, the virus acquired a furin-cleavage site, allowing its S1/S2 boundary to be cleaved in virus-producing cells. In contrast, the S1/S2 boundary of SARS-CoV-1, and indeed all SARS-like viruses isolated from bats, lack this polybasic site and are cleaved by TMPRSS2 or endosomal cathepsins in the target cells^13–20^. Thus the greater stability we observe with S^G614^ would not be relevant to viruses lacking this site, but it appears to be strongly favored when a furin-cleavage site is present. Therefore, the D614G mutation may have emerged to compensate for this newly acquired furin site.

In summary, we show that an S protein mutation that results in more transmissible SARS-CoV-2 also limits shedding of the S1 domain and increases S-protein incorporation into the virion. Further studies will be necessary to determine the impact of this change on the nature and severity of COVID-19.

## Acknowledgements

This work is supported by an administrative supplement to NIH R01 AI129868 for coronavirus research (awarded to M.F. and H.C.)

## Author contributions

H.C. and M.F. conceived of and supervised the study. L.Z., C.B.J., H.M., M.F., and H.C. designed experiments. L.Z., C.B.J., H.M., and A.O. performed experiments. L.Z., C.B.J., H.M., and A.O. analyzed data. E.B.R. and T.I. provided essential reagents. M.F. and H.C. wrote the manuscript. All authors reviewed and edited the manuscript.

## Competing interests

The authors claim no competing interest.

## Data availability

The data that support the findings of this study are available from the corresponding author upon reasonable request.

## Methodsb

### SARS-CoV-2 S protein sequences analysis

To track D614G variation among SARS-CoV-2 isolates, S protein sequences were downloaded from GenBank and separated by the month. Genotype frequency at residue 614 was calculated using R (R Foundation for Statistical Computing) with the Biostrings package. Logo plots of D614G variation were generated by WebLogo after sequence alignment. Total number of sequences analyzed for each month is indicated in the Fig. 1 legend.

### MLV PV production, quantification, and infection

MLV PVs were produced by transfecting HEK293T cells at ~60% confluency in T175 flasks using the calcium-phosphate method with 70 μg of total DNA. The ratio of 5:5:1 by mass was used for the retroviral vector pQCXIX encoding green fluorescence protein (GFP) or firefly luciferase (FLuc), a plasmid expressing MLV gag and pol proteins, and a plasmid expressing the S protein of SARS-CoV-2 or VSV G protein. The plasmid expressing VSV G protein was previously reported^31^. SARS-CoV-2 S protein gene used in the production of MLV PVs was codon-optimized and synthesized by Integrated DNA Technologies based on the protein sequence (GenBank YP_009724390). The S protein gene is fused to the Flag tag sequence (DYKDDDDK) either at its C-terminus or at both the N- and C-termini, as indicated in each experiment. PV-containing culture supernatants were collected at 43 h post transfection, cleared through 0.45 μm filters, and either purified or aliquoted and frozen at −80°C immediately.

PV titers were quantified by RT-qPCR, using primers and a probe that target the CMV promoter. Sense primer: 5’-TCACGGGGATTTCCAAGTCTC-3’, anti-sense primer: 5’-AATGGGGCGGAGTTGTTACGAC-3’, probe: 5’-AAACAAACTCCCATTGACGTCA-3’. Viral RNA was extracted with Trizol (Life Technologies) and GlycoBlue coprecipitant (Invitrogen) and reverse transcribed using the High-Capacity cDNA Reverse Transcription Kit (Applied Biosystems). qPCR was performed using Luna Universal Probe qPCR Master Mix (New England Biolabs) with the known quantity of pQCXIX vector to generate standard curves.

Infection assays were performed by spinoculating (at 2,100 x g for 30 min at 10°C) PVs onto the Mock- and hACE2-293T cells seeded on multiwell plates. Spinoculated plates were incubated for 2 h in a CO2 incubator and medium was replaced with DMEM containing 10% FBS. Infection levels were assessed 24 h post infection by measuring GFP expression by Accuri flow cytometer or luciferase activity using the Luc-Pair Firefly Luciferase HS Assay Kit (GeneCopoeia).

### Analyses of the S protein incorporated into MLV PV

For the analyses of the S protein in the virion, PVs were partially purified. 9 ml of cleared culture supernatants containing PVs (20 ml/T175) were loaded onto 2 ml of 20% sucrose in PBS and centrifuged at 30,000 rpm in the SW41 rotor for 2 h at 10°C^32^. PV pellets were resuspended in 30-50 μl NT buffer (120 mM NaCl, 20 mM Tris, pH8.0). Purified PVs were immediately used or aliquoted and frozen at −80°C. For western blot analyses, 5-10 μl of purified PV, which is equivalent to 0.5-1.0 x 10^10^ vector genomes, was loaded per lane of the 4-12% Bis-Tris gel (Life Technologies), transferred to the PVDF membrane, and blotted with 1 μg/ml anti-Flag M2 antibody (Sigma-Aldrich, F1804) to detect the S-protein bands. 1 μg/ml anti-p30 MLV gag antibody (Abcam, ab130757) and 1:10,000 dilution of goat-anti-mouse IgG-HRP polyclonal antibody (Jackson ImmunoResearch, 115-036-062) were used to detect MLV gag protein as an internal control. Band intensities were measured, using Image Lab software (Bio-Rad). To increase the accuracy of this measurement, the same blots were analyzed several times at different exposures. For silver staining, 30 μl of purified PVs were separated by the 4-12% Bis-Tris gel and stained with Silver Stain Plus (Bio-Rad).

### VLP production and S-protein analysis

SARS-CoV-2 VLPs were produced by transfecting HEK293T cells at ~60% confluency in T75 flasks with 25 μg total DNA using the calcium phosphate method as previously described with a modification^34^. The plasmids expressing SARS-CoV-2 M, N, E, and S proteins were transfected at a ratio of 1:5:5:1. The codon-optimized M, N, and E protein genes were synthesized based on the GenBank protein sequences, YP_009724393, YP_009724397, and YP_009724392, respectively. VLPs were harvested 43 h post transfection from the culture supernatants, cleared by 0.45 μm filtration, and partially purified by pelleting through a 20% sucrose layer as were MLV PVs. VLP pellets were resuspended in 30 μl of NT buffer, and the entire amount was loaded on the 4-12% Bis-Tris gel for WB analyses. As with MLV PV, the S-protein bands were visualized using the anti-Flag M2 antibody, and the N-protein band was detected using pooled convalescent plasma at a 1:500 dilution and 10 ng/ml goat-anti-human IgG antibody conjugated with polymerized HRP (Fitzgerald, 61R-I166AHRP40).

### Cell-surface expression and analysis of the S protein

HEK293T cells, approximately 80% confluent in 6-well plates were transfected with 8 μl PEI 40,000 (Polysciences) and 2 μg plasmid expressing the indicated S protein variant. For ACE2-NN-Ig binding experiments, the S protein constructs with the Flag tag only at the C-terminus, and for the measurement of total S1, those with N-terminal Myc tag (EQKLISEEDL) and C-terminal Flag tag (DYKDDDDK) are used. All tags are fused with a two-glycine (GG) linker. To measure ACE2-binding ability or the level of the S1 domain present on the cell surface, cells were detached two days post transfection with Accutase (Stemcell Technologies Inc.) and incubated with either 1 μg/ml purified hACE2-NN-Ig or anti-Myc antibody (clone 9E10, National Cell Culture Center, Minneapolis, MN), respectively, on ice. hACE2-NN-Ig was previous described and was purified using Protein A-Sepharose CL-4B (GE Healthcare)^31^. To measure the total level of S protein, cells were permeabilized with PBS including 0.5% Triton X-100 (Sigma-Aldrich) at room temperature for 10 min and incubated with 1 μg/ml anti-Flag M2 antibody (Sigma-Aldrich).

### Neutralization assay with human immune plasma

Deidentified blood samples were obtained by the Allergy, Asthma and Immunology Specialists of South Florida, LLC for COVID-19 serotyping, and exempt from human subject research under 45 CFR 45.101(b)(4). MLV PVs encoding firefly luciferase and pseudotyped with the indicated SARS-CoV-2 S protein were preincubated for 1 h at 37°C with or without convalescent or healthy plasma serially diluted in DMEM containing 10% FBS. Mock- and hACE2-HEK293T cells on 96-well plates were infected with the preincubation mixes and infection levels were assessed 24 h later by measuring luciferase activity using the Luc-Pair Firefly Luciferase HS Assay Kit (GeneCopoeia).

### Reagent availability

Plasmids and cell lines used in this study are available from the corresponding author upon request.

### Statistical analysis

All appropriate data were analyzed with GraphPad Prism 7 (GraphPad Software Inc.). All hypothesis tests were performed as two-tailed tests. Specific statistical analysis methods are described in the figure legends where results are presented. Values were considered statistically significant for *p* values below 0.05. Exact *p* values are provided in appropriate figures.

---

**Extended Data Fig. 1.**
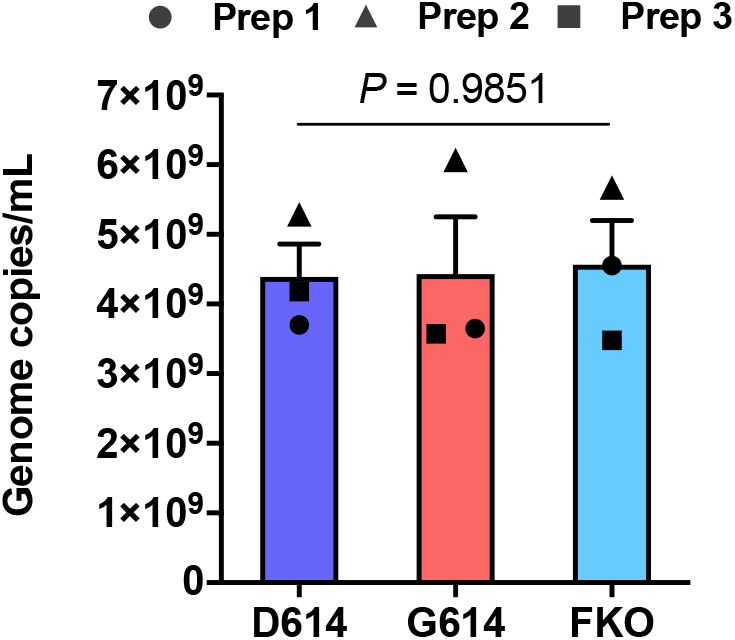
Titers of various MLV PVs. The titers of the MLV PVs used in the experiments shown in Fig. 1c,d were quantified by RT-qPCR. The same symbols in different PV groups indicate they were from the same batch. Each symbol indicates an average value of a duplicated experiment. Similar PV titers validate that enhanced infectivity of PV^G614^ was not resulted from a large difference in virus titers and/or normalization thereof. Mean ± SEM of two independent PV preparations are shown. Significant differences were analyzed by one-way ANOVA.

**Extended Data Fig. 2.**
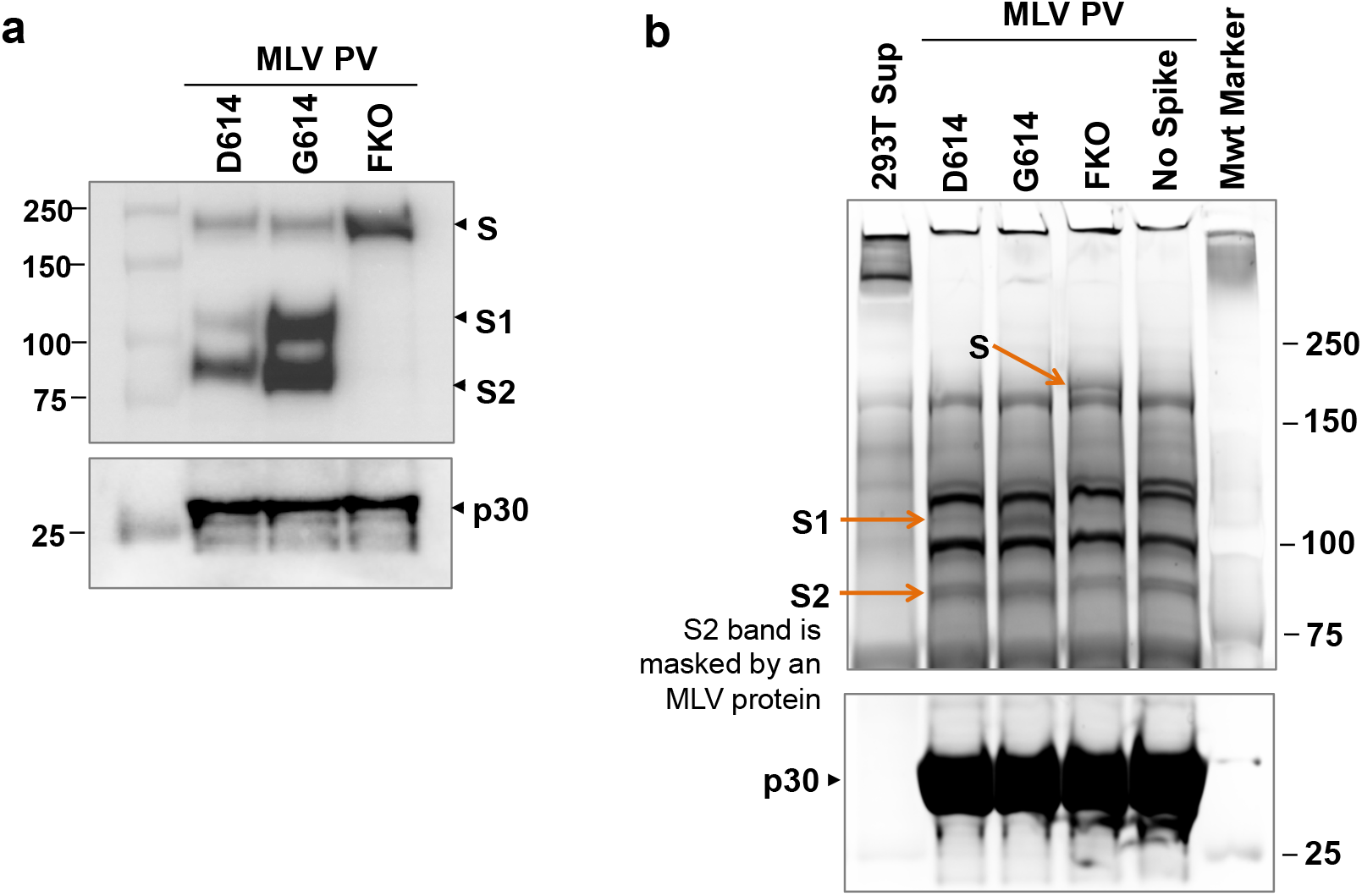
Superior infectivity of G614 results from decreased S1 shedding and higher level of S protein in the virion. **a,** A similar experiment as that shown in Fig. 2c but performed with an independently prepared PV batch. **b,** The same PVs used in the experiment shown in Fig. 2c were analyzed by SDS-PAGE and silver stain to avoid a potential bias caused by using the M2 antibody that recognizes N- and C-terminal Flag tags with different affinity. Although the S2 band is masked by an MLV-derived protein, the S1 band is clearly separated and much weaker in PV^D614^ compared to that in PV^G614^

**Extended Data Fig. 3.**
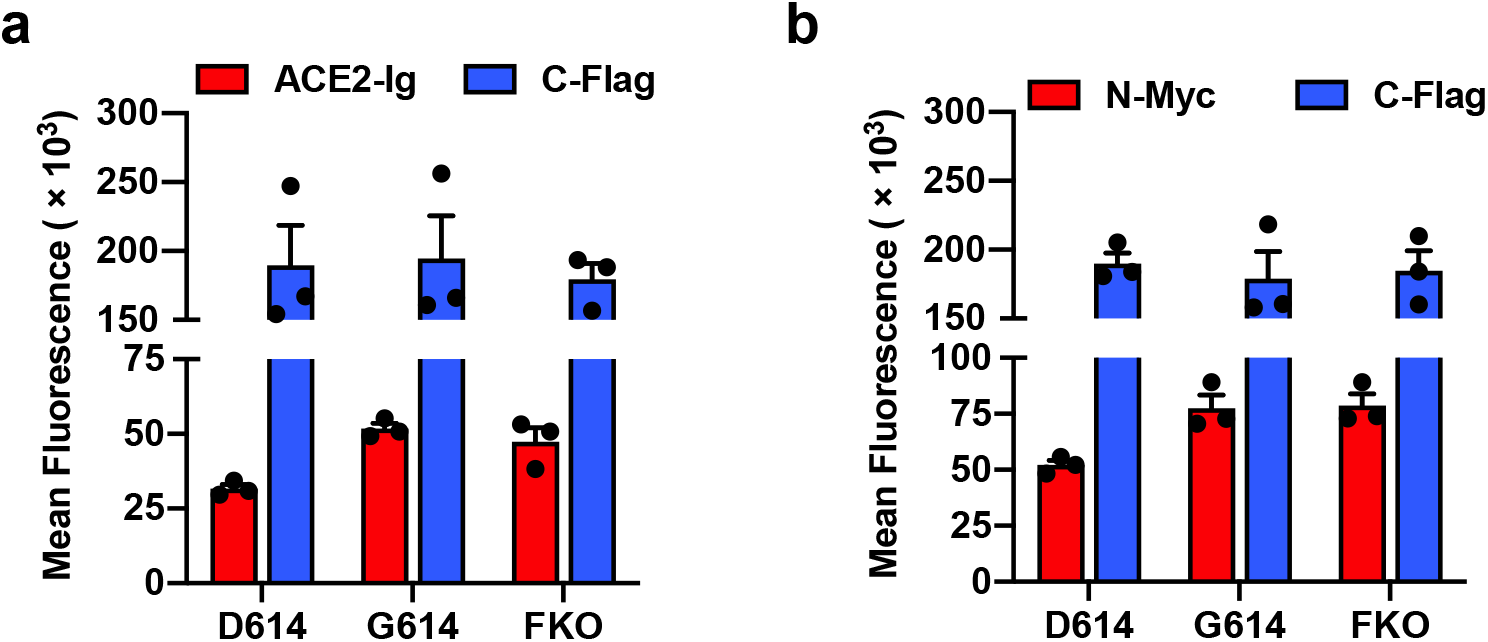
S^G614^ binding to hACE2 is not increased. The same data presented in Fig. 4a and b before normalized to total S protein level. **a,** The S protein containing C-terminal Flag tag is transfected into HEK293T cells and assessed for hACE2-NN-Ig binding. Total S protein was measured by detecting the Flag tag in the permeabilized cells. **b,** Experiments similar to those in (a) except the S protein contains N-Myc and C-Flag tags, and S1 level was assessed using an anti-Myc antibody. Each symbol in (a,b) indicates an average value of a duplicated experiment. Mean ± SEM of three independent experiments are presented.

## Notes

### Competing Interest Statement

The authors have declared no competing interest.

